# PA28γ promotes the malignant progression of tumor by elevating mitochondrial function via C1QBP

**DOI:** 10.1101/2024.07.23.604769

**Authors:** Jiongke Wang, Yujie Shi, Ying Wang, Yingqiang Shen, Huan Liu, Silu Sun, Yimei Wang, Xikun Zhou, Yu Zhou, Xin Zeng, Jing Li, Qianming Chen

**Author notes:** Correspondence and requests for materials should be addressed to: Jing Li, State Key Laboratory of Oral Diseases & National Center for Stomatology & National Clinical Research Center for Oral Diseases & Research Unit of Oral Carcinogenesis and Management & Chinese Academy of Medical Sciences, West China Hospital of Stomatology, Sichuan University, Chengdu 610041, Sichuan, China, Xin Zeng, State Key Laboratory of Oral Diseases & National Center for Stomatology & National Clinical Research Center for Oral Diseases & Research Unit of Oral Carcinogenesis and Management & Chinese Academy of Medical Sciences, West China Hospital of Stomatology, Sichuan University, Chengdu 610041, Sichuan, China. These authors contributed equally to this article (Jiongke Wang, Yujie Shi).

## Abstract

Proteasome activator 28γ (PA28γ) plays a critical role in malignant progression of various tumors, however, its role and regulation are not well understood. Here, using oral squamous cell carcinoma (OSCC) as main research model, we discovered that PA28γ interacted with complement 1q binding protein (C1QBP), which is dependent on the N-terminus of C1QBP rather than the known functional domain (amino acids 168-213). Notably, we found that PA28γ enhances C1QBP protein stability in OSCC. Functionally, PA28γ contributes to the malignant progression of OSCC by affecting mitochondrial morphology and oxidative phosphorylation (OXPHOS) through C1QBP in vitro and vivo. Mechanically, PA28γ upregulates the expression of optic atrophy 1 (OPA1), mitofusin 1 (MFN1), mitofusin 2 (MFN2), and the mitochondrial respiratory complex via C1QBP. Moreover, in a clinical cohort of OSCC patients, PA28γ was positively correlated with C1QBP expression and negatively correlated with prognosis. Therefore, C1QBP represents a potential therapeutic target for cancer treatment and prognosis.

## Introduction

Proteasome activator 28γ (PA28γ), also known as REGγ or PSME3, is a well-known non-ATP-dependent proteasome regulator. PA28γ is overexpressed in several malignant tumors, including thyroid, colon, and breast cancer, and is significantly associated with poor prognosis (Mao, Liu, Li, & Luo, 2008; Stadtmueller & Hill, 2011). By regulating the stability of key components in various complex signaling pathways, PA28γ influences the biological behavior of tumor cells. For instance, PA28γ interacts with both MDM2 and p53 proteins and facilitates their physical interaction, which promotes ubiquitination- and MDM2-dependent proteasomal degradation of p53 and inhibits apoptosis after DNA damage (Zhang & Zhang, 2008). Our previous studies revealed that the expression level of PA28γ in oral squamous cell carcinoma (OSCC) cancer nest tissues is positively correlated with patient prognosis (J. Li et al., 2015; S. Liu et al., 2018). However, its role and regulation are not well understood.

OSCC is the most common oral and maxillofacial malignant tumor(Siegel, Miller, Wagle, & Jemal, 2023; Sung et al., 2021). The pathogenic factors of OSCC include smoking, alcohol consumption, and viral infections, such as human papillomavirus (HPV). Therapies for OSCC include surgery, radiotherapy, chemotherapy and immunotherapy. Despite the rapid development of imaging, surgery, radiotherapy and immunotherapy in recent years, along with the emphasis on personalized treatment and multidisciplinary cooperation in the treatment of OSCC, the overall survival rate of OSCC patients has not significantly improved (Peres et al., 2019). The main reasons for unsatisfactory treatment efficacy include invasion of surrounding tissue, early lymph node metastasis and a tendency toward recurrence (Lindemann, Takahashi, Patel, Osman, & Myers, 2018). Metabolic reprogramming is a crucial characteristic of tumors, closely associated with malignant behaviors such as tumor growth, invasion, metastasis, and immune escape (Ohshima & Morii, 2021). Current research on metabolic reprogramming in OSCC primarily focused on mechanism of glycolytic metabolism and metabolic shift from glycolysis to oxidative phosphorylation (OXPHOS) of oral squamous cell carcinoma, which lays the groundwork for novel therapeutic interventions to counteract OSCC (Chen et al., 2024; Zhang et al., 2020). Understanding the molecular events that lead to the occurrence and development of OSCC can aid in developing new tumor-targeted therapies, which hold significant clinical application potential.

The complement 1q binding protein (C1QBP) is a crucial protein for maintaining mitochondrial function, particularly in mitochondrial OXPHOS (Ghebrehiwet, Geisbrecht, Xu, Savitt, & Peerschke, 2019; Q. Wang et al., 2022). OXPHOS is essential for the production of adenosine triphosphate (ATP), which is necessary for tumor development. C1QBP has been shown to play a significant role in cancer progression, as influencing tumor growth, invasion and metastasis (Bai et al., 2019; Hou, Lu, Wang, & Yang, 2022; Vendramin et al., 2018). Due to these roles, C1QBP presents a promising therapeutic target for various tumors, including melanoma, breast cancer, and colorectal cancer (Matsumoto & Bay, 2021). In OSCC, enhanced mitochondrial OXPHOS function might also be linked to malignant tumor progression (Vyas et al., 2021; Xiao et al., 2021; Zhu, Liu, Wu, & Wang, 2021), although the underlying mechanism remain unclear.

In this study, we discovered that PA28γ interacts with C1QBP and that PA28γ can stabilize C1QBP. This interaction enhances OXPHOS and promotes the growth, migration and invasion of OSCC cells. Additionally, the expression of PA28γ and C1QBP is increased and positively correlated in OSCC. Therefore, PA28γ and C1QBP are potential targets for the treatment and prognosis of cancer. In our knowledge, it is the first study to describe the undiscovered role of PA28γ in promoting the malignant progression of OSCC by elevating mitochondrial function, providing new clinical insights for the treatment of OSCC.

## Results

### PA28γ interacts with and stabilizes C1QBP

To explore how PA28γ promotes OSCC progression, we conducted a gene□gene interaction analysis using the GeneMANIA database. This analysis identified C1QBP as a significant gene within the interaction network, with the protein-coding gene of PA28γ also playing a key role in the C1QBP interaction network (Appendix Fig. 1A, B). Subsequently, pull-down and coimmunoprecipitation analysis demonstrated that both endogenous and exogenous PA28γ and C1QBP could bind to each other (Fig. 1A, B and Appendix Fig. 1 C, D). Moreover, proximity ligation assay (PLA) (Fig. 1C) revealed a close proximity between PA28γ and C1QBP in cells, thereby suggesting a potential interaction between the two proteins. Then, we developed four OSCC cell lines with stable overexpression of PA28γ using a Flag-PA28γ-expressing lentivirus and observed that the protein level of C1QBP was higher in these cell lines compared to control cells (Fig. 1D). Notably, PA28γ upregulated C1QBP protein levels in a dose-dependent manner (Fig. 1E). To determine whether the upregulated C1QBP was due to increased stability, cells were pretreated with cycloheximide (CHX) to inhibit protein synthesis. As a result, in the presence of PA28γ, C1QBP exhibited a significantly lower turnover rate compared to the control group (Fig. 1F). In addition, PA28γ stabilized C1QBP in the absence and presence of a proteasome inhibitor MG-132 (Appendix Fig. 1E), whereas the inhibitor alone did not stabilize C1QBP. Furthermore, AlphaFold 3-based structural modeling predicted that the PA28γ-C1QBP interaction could be mediated by the N-terminal region of C1QBP (amino acid residues 1-167), rather than the known functional domain (amino acids 168-213) that bind to mitochondrial antiviral proteins (Peerschke et al., 2020; Xu, Xiao, Liu, Ren, & Gu, 2009) (Appendix Fig. 1F-I). Results from a pull-down analysis of the PA28γ interaction with full length and truncated C1QBP proteins are consistent with this prediction (Fig. 1G-I). In addition, we performed energy minimization to refine the structural model for PA28γ-C1QBP complex, and used the refined model to reveal potential interactions between the two proteins. The results suggest that the T76 and G78 residues in C1QBP would form hydrogen bonds with the D177 residue in PA28γ (Appendix Fig. 1J). Consistently, coimmunoprecipitation analysis demonstrated that mutations that would disrupt these hydrogen bonds (T76A and G78N) significantly reduced the binding of C1QBP to PA28γ (Fig. 1J). Therefore, our results illustrated that PA28γ could interacts with C1QBP, in a manner dependent on the N-terminus of C1QBP, and that PA28γ could stabilizes C1QBP.

**Fig. 1.**
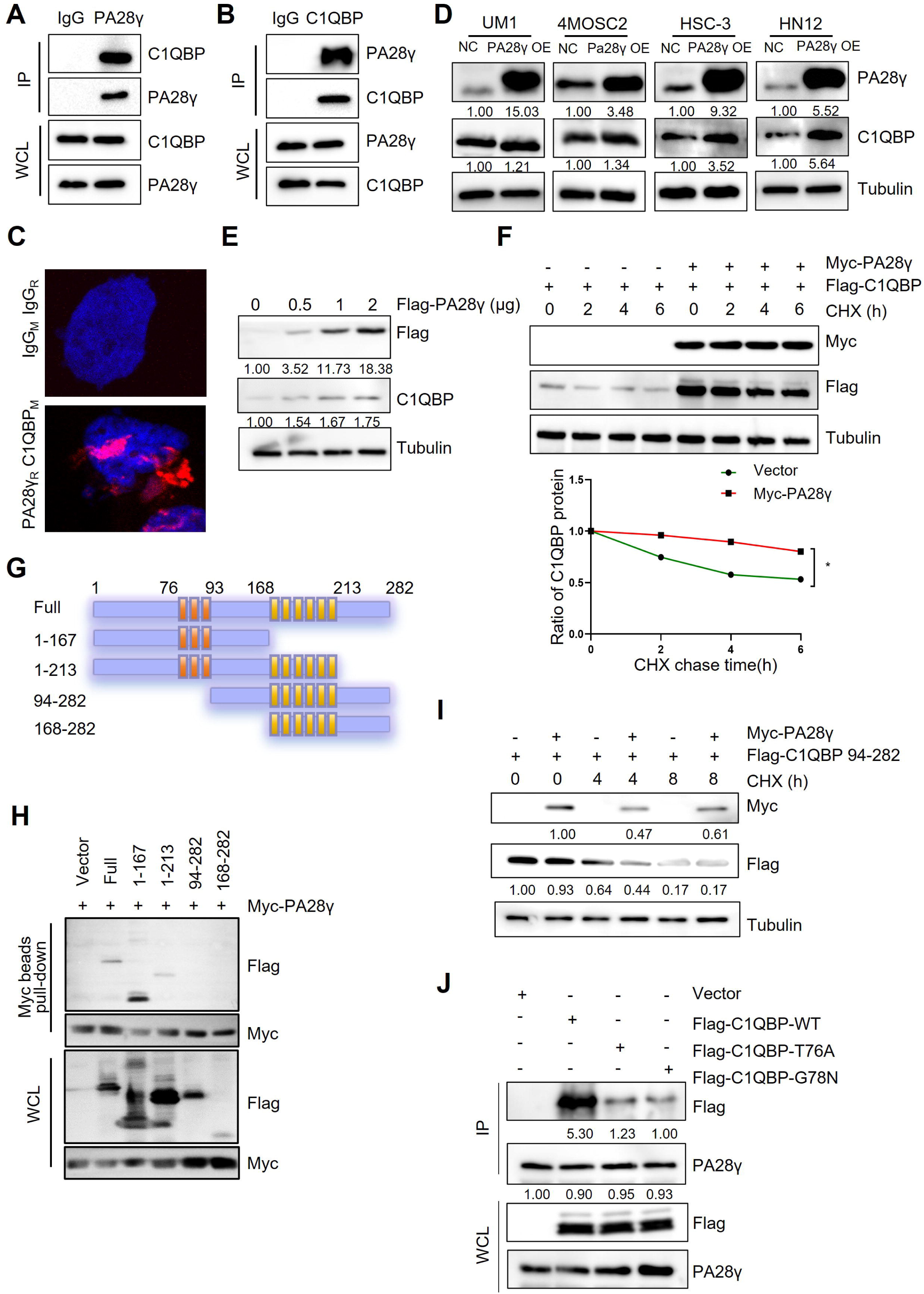
The interaction between PA28γ and C1QBP. **(A, B)** The interaction between endogenous PA28γ and C1QBP in HSC-3 cells was verified via immunoprecipitation (IP). **(C)** PLA image of UM1 cells shows the interaction between C1QBP and PA28γ in both cytoplasm and nucleus (red fluorescence). **(D)** Western blot analysis of C1QBP in four OSCC cell lines with PA28γ-overexpressing. **(E)** Western blot analysis of C1QBP in 293T cells transfected with increasing doses of Flag-PA28γ. **(F)** 293T cells transfected with Flag-C1QBP with or without Myc-PA28γ were treated with CHX (100 μg/ml) for the indicated periods of time. Quantification of Flag-C1QBP levels relative to tubulin levels is shown (the data are representative of 1 experiment with 3 independent biological replicates, \**P*<0.05). **(G)** Full-length C1QBP and truncation with deletion of functional domains. **(H)** Pull-down of 293T cells transfected with Myc-PA28γ and full-length Flag-C1QBP or truncation mutants of functional domains for 36h. **(I)** 293T cells transfected with Flag-C1QBP 94-282 with or without Myc-PA28γ were treated with CHX (100 μg/ml) for the indicated periods of time. **(J)** IP of 293T cells transfected with Flag-C1QBP wild type or mutations (T76A and G78N) for 48h.

### PA28γ and C1QBP colocalize in mitochondria and affect mitochondrial functions

To further investigate the physical interaction between PA28γ and C1QBP, we conducted an immunofluorescence (IF) assay in two OSCC cell lines, as shown these two proteins were colocalized in the mitochondria (Fig. 2A). Remarkably, transmission electron microscopy (TEM) images revealed fewer mitochondrial vacuoles and higher mitochondrial ridge density in PA28γ-overexpressing cells compared to control cells (Fig. 2B and Appendix Fig. 2A). IF analysis also showed increased mitochondrial lengths and areas in the PA28γ-overexpressing cells (Fig. 2C, D). Following that, we conducted a Cell Mito Stress Test to measure the oxygen consumption rate (OCR) (Fig. 2E, F). The results indicated significantly higher basal respiration, maximal OCRs and ATP production in PA28γ-overexpressing cells compared to control cells (Fig. 2G-I and Appendix Fig. 2B-D). Considering that mitochondria are the primary site for ROS generation and that ROS play a crucial role in carcinogenesis, we measured ROS levels in OSCC cells. Consistently, ROS levels were significantly higher in PA28γ-overexpressing cells (Fig. 2J and Appendix Fig. 2E). In addition, experiments conducted in PA28γ-sh OSCC cells yielded opposite results, further confirming these conclusions (Appendix Fig. 2F-K).

**Fig. 2.**
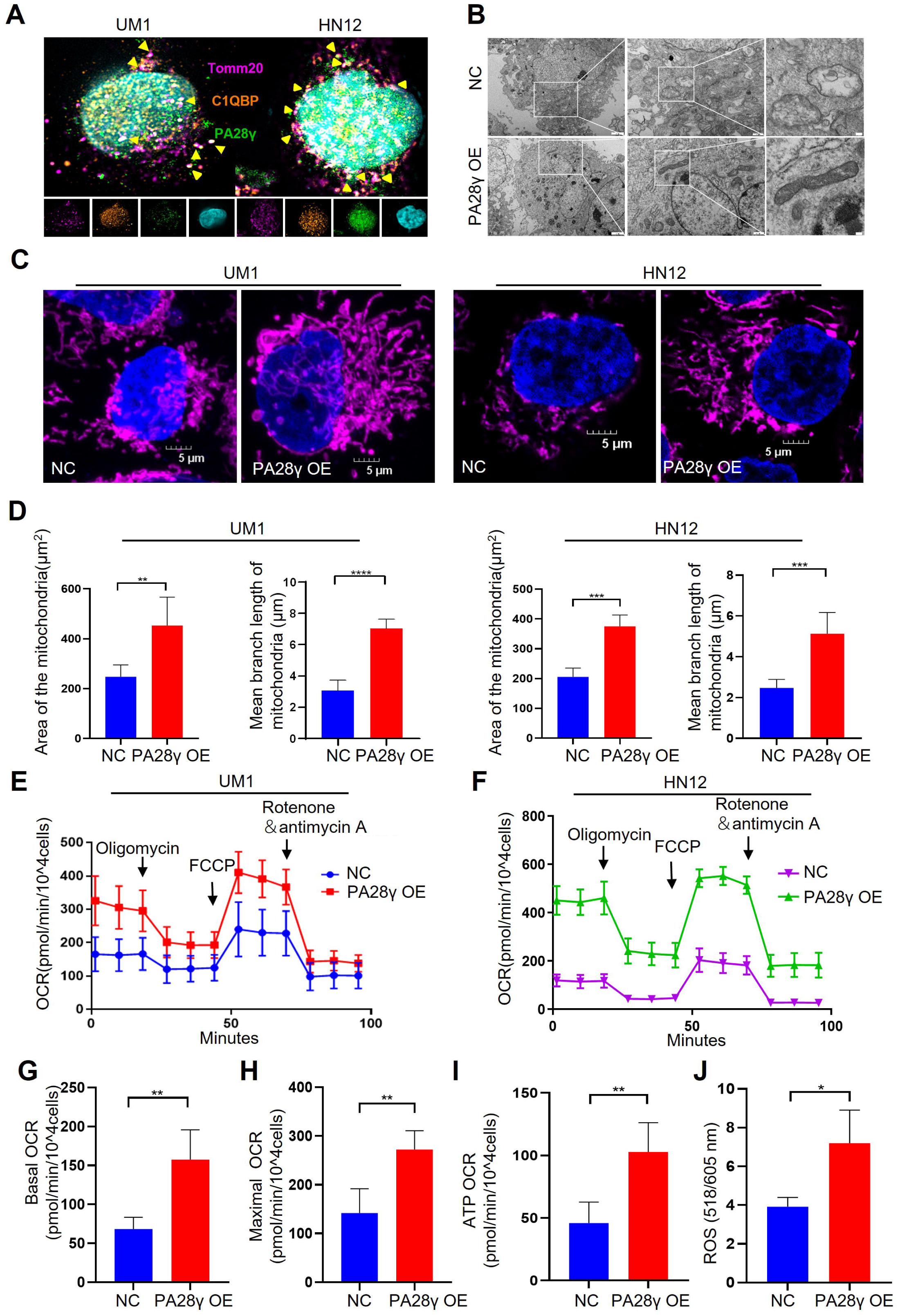
PA28γ and C1QBP colocalize in mitochondria and influence mitochondrial functions *in vitro*. **(A)** Confocal image of IF in two OSCC cell lines. **(B)** TEM images of PA28γ-overexpressing and control UM1 cells. **(C)** Representative confocal images of mitochondria in two OSCC cell lines. **(D)** The area and mean branch length of mitochondria in two OSCC cell lines were measured by ImageJ (the data are presented as the mean ± SD of 3 independent experiments; **P<0.01, ***P<0.001, and ****P<0.0001). **(E, F)** OCRs of PA28γ-overexpressing and control UM1 and HN12 cells were plotted using a Cell Mito Stress Test Kit (the data are presented as the means ± SDs of 3 independent experiments). **(G-I)** Basal OCRs, maximal OCRs and ATP production of PA28γ-overexpressing and control UM1 cells measured by the Cell Mito Stress Test (the data are presented as the means ± SDs of 3 independent experiments, \*\**P*<0.01). **(J)** ROS generation in PA28γ-overexpressing and control UM1 cells (the data are presented as the means ± SDs of 3 independent experiments; * *P*<0.05).

To demonstrate the carcinogenic abilities of PA28γ and C1QBP, we injected 4MOSC2 cells, a mouse OSCC line, with or without Pa28γ overexpression under the surface of the tongue in C57BL/6 mice (Appendix Fig. 3A). After 10 days of normal feeding, the tumors in the Pa28γ-overexpressing groups were significantly larger than those in the control group. Furthermore, via immunohistochemical (IHC) staining revealed that that PA28γ overexpression upregulate C1QBP in vivo (Fig. 3A-C). Consistent with the in vitro experiments, ATP production and ROS generation were significantly increased (Fig. 3D, E). Additionally, analysis of OSCC xenograft tumor tissue in nude mice revealed that PA28γ’s regulation of C1QBP in OSCC cells is independent of the immune system (Appendix Fig. 3B, C). In addition, we established stable Pa28γ-silenced B16 cells, a mouse melanoma cell line, and found that the level of C1qbp protein was decreased in Pa28γ-silenced B16 cells (Appendix Fig. 3D). Cells with or without Pa28γ silencing were grafted into the flanks of Pa28γ ^ko/ko^ and Pa28γ ^wt/wt^ C57BL/6 mice by subcutaneous injection (Appendix Fig. 3E). Remarkably, the tumors derived from the silenced Pa28γ cells in Pa28γ ^ko/ko^ mice were smaller than those in the other groups (Fig. 3F-H). The trend of ATP and ROS was consistent with tumor size (Fig. 3I, J), and the protein levels of C1qbp in the tumors were also reduced in the Pa28γ-silenced groups (Fig. 3K).

**Fig. 3.**
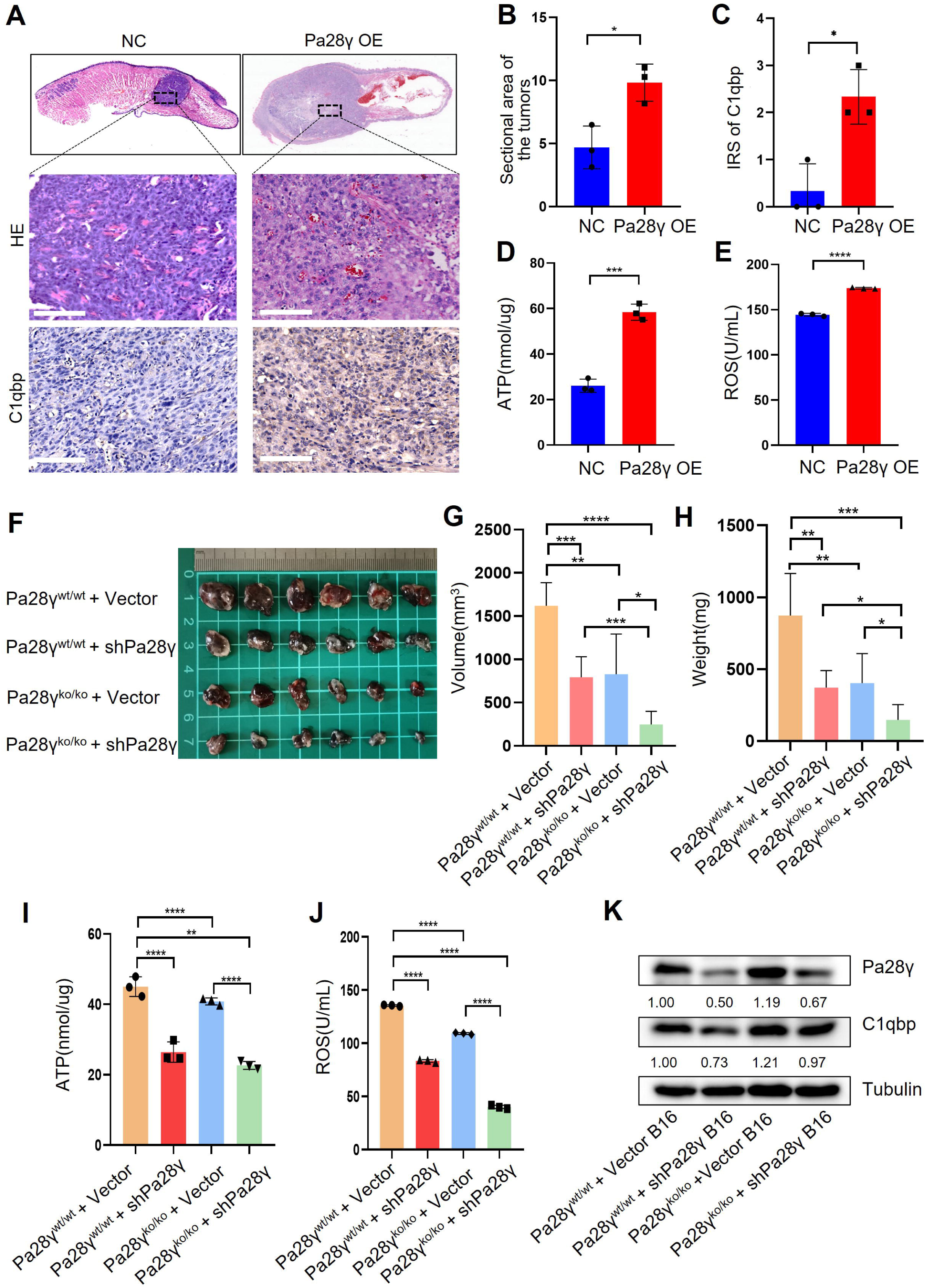
PA28γ affect mitochondrial functions *in vivo*. **(A)** Representative images of H&E staining and IHC staining of C1qbp of tongue sections from mice (n=3). **(B)** Quantification of the sectional area in tumors from the Pa28γ-overexpressing and control groups (n=3; the data are presented as the means ± SDs of 3 samples; *\*P*<0.05). **(C)** Comparison of the immunoreactive scores (IRSs) of C1qbp antibody staining in the Pa28γ-overexpressing and control groups (n=3, the data are presented as the means ± SDs; *P<0.05). **(D, E)** Quantification of ATP production and ROS levels in tumors from the Pa28γ-overexpressing and control groups (n=3; the data are presented as the means ± SDs of 3 samples; *\*P*<0.05, \*\**\*P*<0.001, *\*\*\*\*P*<0.0001). **(F)** Images of the tumors in different groups at the endpoint (n=6). **(G-J)** The volume, weight, ATP and ROS of tumors in different groups at the endpoint (n=6; the data are presented as the means ± SDs of 3 independent experiments; *\*P*<0.05, **P<0.01, \*\**\*P*<0.001, *\*\*\*\*P*<0.0001). **(K)** Western blot analysis of tumors in different groups.

Collectively, these data suggest that PA28γ, which co-localizes with C1QBP in mitochondria, may involve in regulating mitochondrial morphology and function.

### PA28γ regulates mitochondrial OXPHOS by upregulating C1QBP

Due to the results of targeted metabolomics analysis by mass spectrometry (Appendix Fig. 4A, B), it was found that PA28γ may affect the biological behavior and function of OSCC by influencing metabolism. Therefore, we explored the protein levels of the mitochondrial respiratory chain complex, C1QBP and other mitochondrial functional proteins. We found that the levels of C1QBP, complex I and IV proteins, as well as OPA1, MFN1 and MFN2 proteins, were upregulated in PA28γ-overexpressing OSCC cells (Fig. 4A, B and Appendix Fig. 4C, D), suggesting that the effect of PA28γ on the levels of these mitochondrial fusion factors is mediated by C1QBP via a molecular mechanism that is currently unknown. These increases could be reversed by C1QBP silencing in PA28γ-overexpressing OSCC cells (Fig. 4C).

**Fig. 4.**
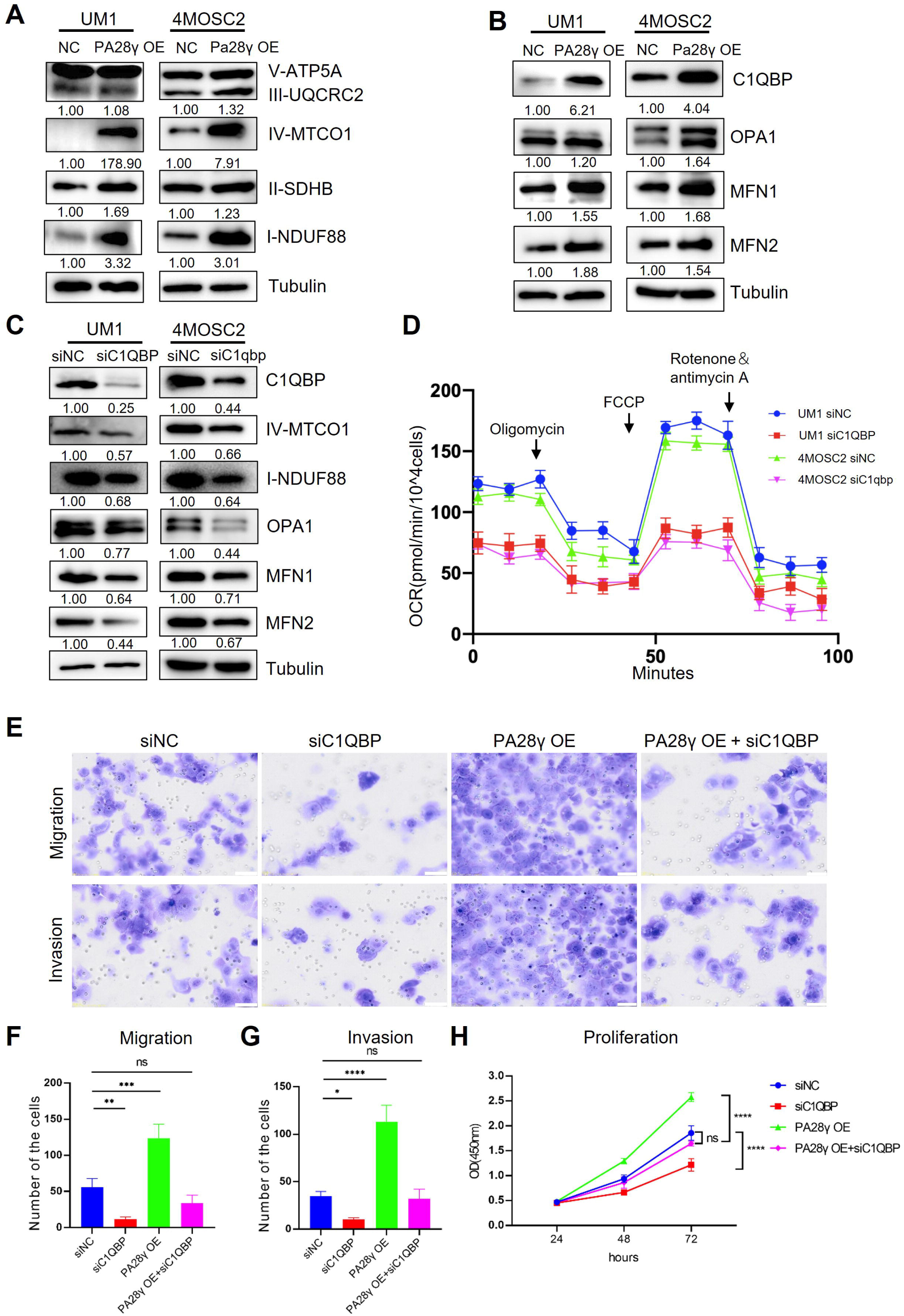
PA28γ regulates mitochondrial OXPHOS and cellular biological behavior via C1QBP. **(A, B)** Western blot analysis of PA28γ-overexpressing and control UM1 and 4MOSC2 cells. **(C)** Western blot analysis of PA28γ-overexpressing OSCC cells transfected with siNC or siC1QBP. **(D)** OCRs of C1QBP-silenced and control PA28γ-overexpressing UM1 and 4MOSC2 cells (the data are presented as the means ± SDs of 3 independent experiments). **(E-H)** Cell migration, invasion and proliferation in control, C1QBP-silenced, PA28γ-overexpressing and PA28γ-overexpressing + C1QBP-silenced UM1 cells (the data are presented as the means ± SDs of 3 independent experiments; \**P*<0.05, \*\**P*<0.01, \*\**\*P*<0.001, *\*\*\*\*P*<0.0001).

Consistent with this, the OCRs of C1QBP-silenced PA28γ-overexpressing OSCC cells were markedly lower than those of control cells (Fig. 4D and Appendix Fig. 4E-G). Subsequently, we constructed a series of experiments to detect the biological behavior of control and PA28γ-overexpressing OSCC cells with or without C1QBP silencing. C1QBP knockdown significantly attenuated the migration, invasive and proliferation capabilities previously augmented by PA28γ overexpression (Fig. 4E-H). These data suggest that PA28γ enhances mitochondrial OXPHOS function through C1QBP.

### C1QBP expression was positively associated with PA28γ expression and was associated with a worse prognosis in patients with OSCC or SKCM

To determine the correlation between PA28γ and C1QBP, IHC staining was performed in the clinical cohort of oral mucosa carcinogenesis. The results showed that both C1QBP and PA28γ were notably upregulated during the progression of oral mucosa carcinogenesis (Fig. 5A, B and Appendix Fig. 5A), and the levels of PA28γ and C1QBP were positively related (Fig. 5C). Similarly, both C1QBP and PA28γ were upregulated in metastatic OSCC tissues (Fig. 5D, E and Appendix Fig. 5B), and the levels of PA28γ and C1QBP were consistently positively correlated in OSCC cohort (Fig. 5F). Notably, consistent with our previous findings on PA28γ (J. Li et al., 2015), C1QBP or combining PA28γ and C1QBP could be a negative predator in our multicenter OSCC clinical cohort (Fig. 5G, H), TCGA HNSC database (Fig. 5I, J) and TCGA SKCM database (Appendix Fig. 5C, D).

**Fig. 5.**
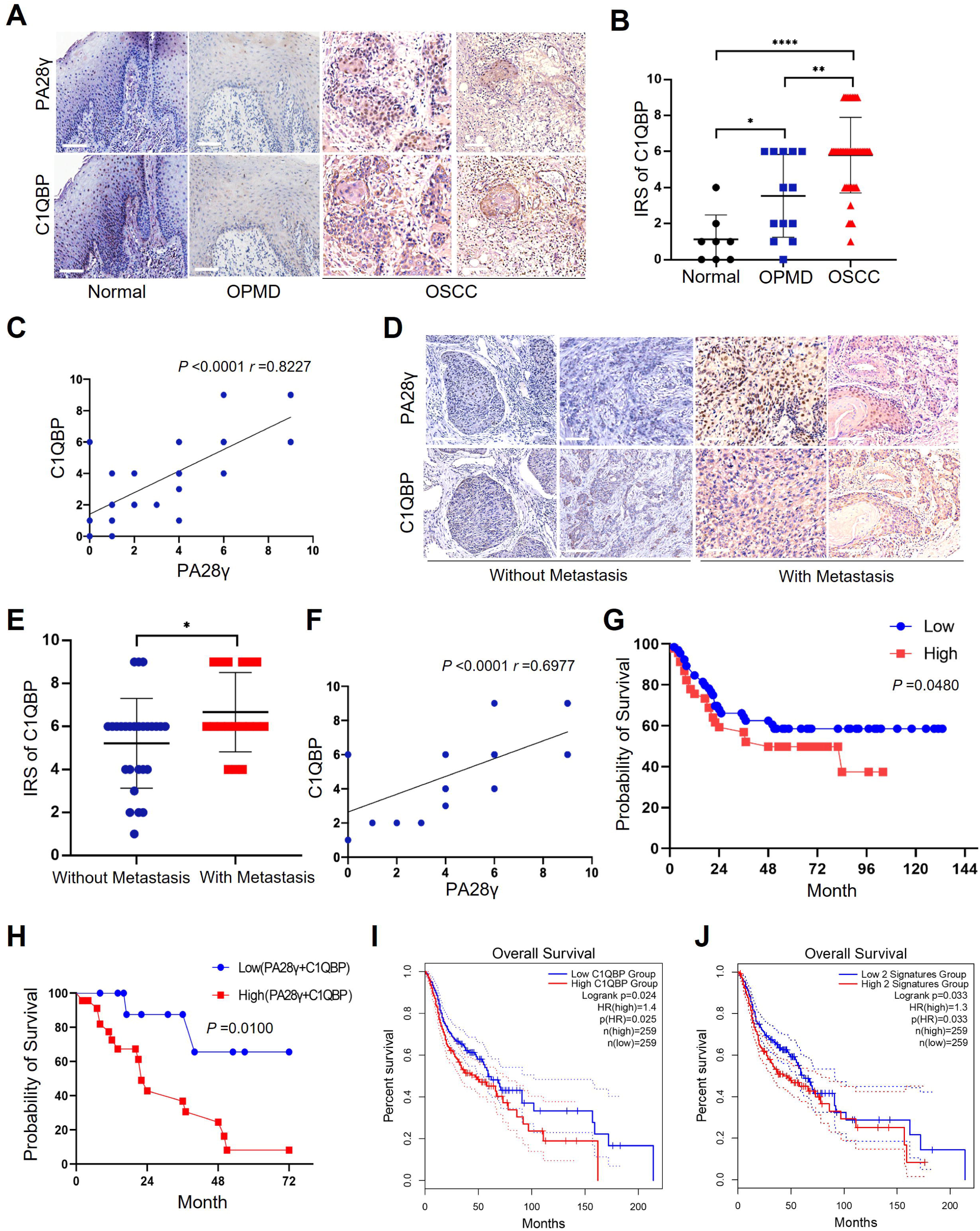
The correlation between PA28γ and C1QBP in the carcinogenesis and development of OSCC. **(A)** Representative IHC staining of PA28γ and C1QBP in normal (n=8), OPMD (n=13) and OSCC (n=45) samples. **(B)** Comparison of the immunoreactive scores (IRSs) of C1QBP between the normal, OPMD and OSCC groups (the data are presented as the means ± SDs; *\*P*<0.05, \**\*P*<0.01, *\*\*\*\*P*<0.0001). **(C)** Spearman correlation analysis was used to test the correlation between PA28γ and C1QBP in normal, OPMD and OSCC tissues (*P*<0.0001, *r*=0.8227). **(D)** Representative IHC staining of PA28γ and C1QBP in nonmetastatic (n=27) and metastatic (n=18) OSCC patients. **(E)** Comparison of the IRSs of C1QBP in the nonmetastatic and metastatic OSCC groups (the data are presented as the means ± SDs; *\*P*<0.05). **(F)** Spearman correlation analysis was used to test the correlation between PA28γ and C1QBP in OSCC tissues (*P*<0.0001, *r*=0.6977). **(G)** Kaplan–Meier analysis of the protein expression of C1QBP in our multicenter OSCC clinical cohort (n=295, *P*=0.0480). **(H)** Kaplan–Meier analysis of both low or high protein expression of C1QBP and PA28γ in our multicenter OSCC clinical cohort (n=295, *P*=0.0100). **(I)** Kaplan–Meier analysis of the protein expression of C1QBP in TCGA HNSC database (n=259, *P*=0.025). **(J)** Kaplan–Meier analysis of both low or high protein expression of C1QBP and PA28γ in TCGA HNSC database (n=259, *P*=0.033).

## Discussion

Metabolic reprogramming is one of the hallmarks of malignant tumors and is related to the malignant biological behavior of tumors (Faubert, Solmonson, & DeBerardinis, 2020; Hanahan, 2022; Tsai et al., 2023; Xia et al., 2021). The metabolic phenotype of tumor cells differs from that of normal cells and dynamically changes during tumor progression. OSCC is a common head and neck malignancy that is still a growing global health problem (Johnson et al., 2020). The development of OSCC is a complex and multifaceted process in which metabolic reprogramming appears to be important (B. Liu, Si, Wei, Zhang, & Chen, 2023). However, the mechanism of metabolic reprogramming remains unclear. Excitingly, we found the evidence that PA28γ interacts with and stabilizes C1QBP. We speculate that aberrantly accumulated C1QBP enhances the function of mitochondrial OXPHOS and leads to the production of additional ATP and ROS by activating the expression and function of OPA1, MNF1, MFN2 and mitochondrial respiratory chain complex proteins. This process results in resulting in mitochondrial fusion and malignant tumor progression (Fig. 6). Our study highlights the underlying mechanism by which PA28γ participates in the regulation of OXPHOS by upregulating C1QBP in OSCC.

**Fig. 6.**
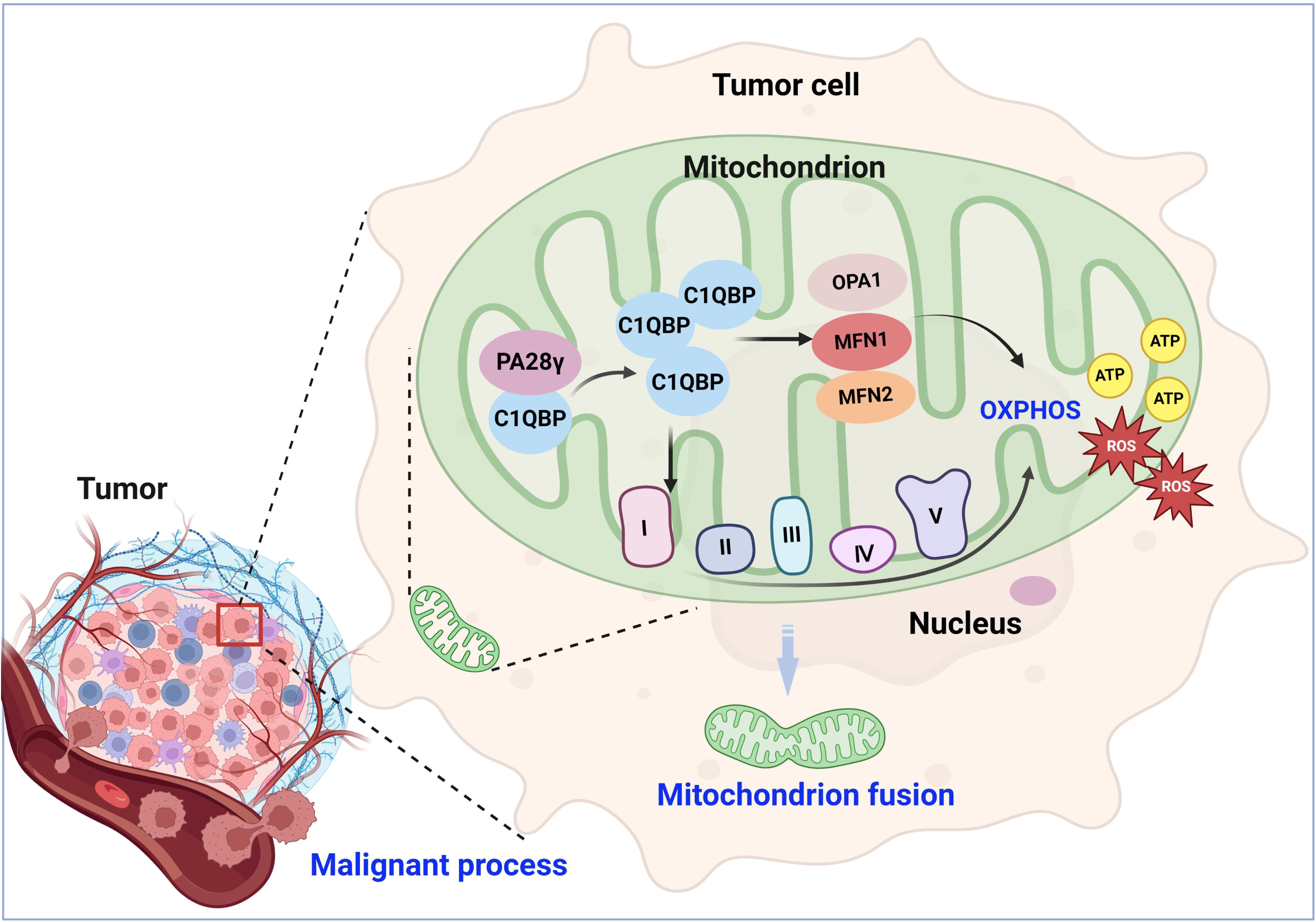
Molecular mechanism through which PA28γ interacts with C1QBP in the malignant progression of tumor. PA28γ can interacts with and stabilize C1QBP, which can activate the expression and function of OPA1, MNF1, MFN2 and mitochondrial respiratory chain complex proteins, resulting in enhanced mitochondrial OXPHOS, mitochondrial fusion and malignant tumor progression.

C1QBP is also called gClqR, p32, p33 and HABP1, can be distributed inside cells, on the cell surface, and can also be secreted extracellularly (Egusquiza-Alvarez & Robles-Flores, 2022). Within cells, C1QBP is primarily located in the mitochondria. The key functions of mitochondrial C1QBP include maintaining OXPHOS, supplying energy to cells, and achieving homeostasis of mitochondrial metabolism (Ghebrehiwet et al., 2019; Q. Wang et al., 2022). C1QBP is highly expressed in cells that require a large amount of energy, such as the heart, skeletal muscle, testis, ovary, small intestine and colon (Feichtinger et al., 2017). Sufficient energy is essential for cancer development, and C1QBP promotes malignant behaviors such as tumor invasion and metastasis by enhancing OXPHOS function in various cancers (Raschdorf et al., 2021; Q. Wang et al., 2022). However, there are no reports on the expression and functions of C1QBP in OSCC. In our study, we found the evidence that PA28γ interacts with and stabilizes C1QBP. As a proteasome activator, PA28γ usually affects the development of tumors by regulating the stability of various important proteins. For example, PA28γ can promote the degradation of p53, p21 leading to cancer progression (J. Liu et al., 2010). In addition, PA28γ can also play as a non-degradome role on tumor angiogenesis. For example, PA28γ can regulate the activation of NF-κB to promote the secretion of IL-6 and CCL2 in OSCC cells, thus promoting the angiogenesis of endothelial cells ( S. Liu et al., 2018). However, the function of mitochondrial PA28γ may be differ from that of nuclear PA28γ (Mao et al., 2008). Our study reveals that PA28γ interacts with C1QBP and stabilizes C1QBP at the protein level. Therefore, we speculate that the binding sites of PA28γ and C1QBP may mask the specific post-translational modification sites of C1QBP and inhibit its degradation. For example, the transcriptional coactivator p300 interacts with Smad7 by acetylating two lysine residues in its N-terminus, which stabilize Smad7 and protect it from TGFβ-induced degradation (Grönroos, Hellman, Heldin, & Ericsson, 2002). Overall, the specific mechanism involved in the interaction between PA28γ and C1QBP still needs further exploration.

Considering that C1QBP is a vital protein for maintaining mitochondrial metabolism (Tian et al., 2023), we further investigated the function underlying the malignant progression of OSCC through the interaction between PA28γ and C1QBP in vitro and in vivo. PA28γ and C1QBP colocalized in the mitochondria, and the stabilization of C1QBP by PA28γ enhanced mitochondrial function and OXPHOS. This enhancement is crucial for ATP production, and ATP is the primary energy currency of cellular metabolism. Therefore, PA28γ’s regulation of OXPHOS may impact cellular energy metabolism. PA28γ has been reported to activate the mTORC1 signaling pathway in hepatocellular carcinoma cells to promote glycolysis and inhibit OXPHOS (Yao et al., 2021). These findings contrast with our study, potentially due to organ heterogeneity or the complexity of metabolic reprogramming.

In the tissue microenvironment of oral lichen planus, a potentially malignant oral disorder, PA28γ in epithelial cells can regulate T cell differentiation (Y. Wang et al., 2024), while PA28γ in CAFs is involved in the crosstalk between stromal cells and tumor cells (Z. Li et al., 2024). This suggests that PA28γ may interact with the tumor immune microenvironment. Our phenotypically similar OSCC xenograft tumor models in both immunocompetent and nude mice indicate that the regulation of mitochondrial OXPHOS by PA28γ localized in tumor cell mitochondria does not entirely depend on the immune system (Z. Wang et al., 2019).

Furthermore, our study reveals that PA28γ can regulate C1QBP and influence mitochondrial morphology and function by enhancing the expression of OPA1, MFN1, MFN2 and the mitochondrial respiratory complex. Mitochondrial fusion, crucial for oxidative metabolism and cell proliferation, is regulated by MFN1, MFN2, and OPA1. The first two fuse with the outer mitochondrial membrane, while the last fuses with the inner mitochondrial membrane (Westermann, 2010). OPA1 is an essential inner mitochondrial membrane protein with multiple functions, including regulating mitochondrial fusion, cristae morphology, mtDNA stability, and interacting with the mitochondrial respiratory chain to regulate OXPHOS (Del Dotto et al., 2018).

Finally, we analyzed the expression patterns and significance of PA28γ and C1QBP in a clinical cohort. PA28γ and C1QBP were negatively correlated with prognosis in the OSCC clinical cohort. Notably, some studies have shown that OXPHOS is upregulated in various tumors, with some advanced tumors preferring OXPHOS metabolically (Ohshima & Morii, 2021; Qiu, Li, & Zhang, 2023). OXPHOS activity is associated with the recurrence of OSCC (Noh et al., 2023). Additionally, the enhancement of OXPHOS and ATP production by ROS proto-oncogene 1, which localizes to mitochondria in OSCC, promotes the OSCC invasion(Noh et al., 2023; Grimm et al., 2014). These findings are consistent with our results. Our study not only explain the mechanisms and signaling networks underlying the oncogenic potential of PA28γ but also contribute to understanding the possible mechanisms involved in OSCC metabolic reprogramming.

In summary, we found that PA28γ could interact with C1QBP and stabilize C1QBP. This enhances the function of OXPHOS, leading to increased production of ATP and ROS by increasing the expression and function of OPA1, MNF1, MFN2 and the mitochondrial respiratory chain complex. This process promotes mitochondrial fusion and malignant tumor progression. The expression of PA28γ and C1QBP was increased and positively correlated in OSCC. High expression of PA28γ and C1QBP was correlated with poor prognosis in OSCC patients, indicating that both proteins, in particular their molecular interactions, could serve as potential targets for the treatment and prognosis of OSCC.

## Materials and methods

### Patients and Follow-up

Our study included 4 cohorts. The first cohort included 8 normal controls, 13 patients with oral potentially malignant disorder (OPMD), and 45 patients with OSCC (Appendix Table 1). These patients visited the West China Hospital of Stomatology in 2021. The normal tissues were obtained from patients who underwent maxillofacial plastic surgery. The second cohort for survival analysis included 295 patients diagnosed with primary OSCC tumors who visited West China Hospital of Stomatology, Peking University Hospital of Stomatology and Guangdong Provincial Stomatological Hospital from 2005 to 2009 (Appendix Table 2). None of the patients had cancer in other organs. These patients received regular follow-up for 1-133 months after the operation. The protocol of this study was approved by the ethics committee of the West China Hospital of Stomatology (Approval number: WCHSIRB-D-2020-046). The third cohort, comprising 518 patients with head and neck squamous cell carcinoma (HNSCC), was obtained from The Cancer Genome Atlas (TCGA) database (Appendix Table 3). The last cohort, comprising 458 patients with skin cutaneous melanoma (SKCM), was obtained from TCGA database (Appendix Table 4).

### Laboratory Experiments

All animal studies were approved by the Animal Care and Use Committee for the State Key Laboratory of Oral Diseases (Approval number: WCHSIRB-D-2022-032) in compliance with the Guide for the updated Animal Research: Reporting of In Vivo Experiments (ARRIVE) 2.0 guidelines. Other methods are detailed in the *Supplemental Materials and Methods*, including cell culture and cell-related experiments, animal models, mitochondrial function detection, western blot/immunoprecipitation assays, hematoxylin and eosin (H&E) staining, and immunohistochemistry.

### Statistical Analysis

The data were analyzed using GraphPad Prism software version 8.0. The statistical data are expressed as the mean ± standard deviation (SD). Chi-square and Fisher exact tests, Spearman correlation tests, Wilcoxon rank-sum tests, one-way ANOVA and Student’s t tests were used to analyze the data. The Kaplan□Meier method was used for survival analysis, and the log-rank test was used to evaluate the prognostic value of C1QBP in OSCC patients. The data are shown as the mean ± standard error, and a value of *P*<0.05 was considered to indicate statistical significance: *P*<0.05*, *P*<0.01**, *P*<0.001***, and *P*<0.0001****. Each experiment was repeated 3 times.

## Acknowledgments

This work was supported by grants from the National Natural Science Foundation of China (82072999 and 82273320 to J. Li, 82270986 and 82470983 to X. Zeng, 82201074 to J. Wang), and the Province Natural Science Foundation of Sichuan (23NSFSC0145 to J. Li). The authors thank Zhijun Sun (School and Hospital of Stomatology, Wuhan University) for providing the cells needed for the experiment and Ning Ji (State Key Laboratory of Oral Diseases, Sichuan University) for excellent assistance with microscopic imaging.

## Author contributions

J. Wang, Y. Shi, X. Zeng, J. Li contributed to conception and design, data acquisition, analysis, and interpretation, drafted and critically revised the manuscript; Y. Wang, Y. Shen, H Liu contributed to data acquisition; S. Sun, Y. Wang contributed to data analysis and interpretation; X. Zhou, Q. Chen contributed to conception and design, critically revised the manuscript. All the authors gave final approval and agreed to be accountable for all aspects of the work.

## Conflict of interests

The authors declared no potential conflicts of interest with respect to the research, authorship, and/or publication of this article.

## Notes

### Competing Interest Statement

The authors have declared no competing interest.

### Summary of Updates

Modified some of the textual descriptions in the manuscript

