## Supplementary material for "PA28γ promotes the malignant progression of tumor by elevating mitochondrial function via C1QBP": NA

**Supplemental Materials and Methods**

***Cell Culture***

UM1 and HSC-3 are tongue squamous cell carcinoma cell lines with different invasion and migration capabilities ^1,2^. UM1 cells are derived from primary tongue squamous cell carcinoma, while HSC-3 and HN12 cells are derived from metastatic cervical lymph nodes. The mouse OSCC line 4MOSC2 (gifted by J. Silvio Gutkind from the University of California, San Diego with Materials Transfer Agreement [MTA]: sd-2017-202) ^3^ was cultured in keratinocyte media (Invitrogen, USA) supplemented with Defined Keratinocyte-SFM Growth Supplement (Invitrogen, USA), 5 ng/mL EGF recombinant mouse protein (Invitrogen, USA) and cholera toxin (Sigma, USA) at 37°C and 5% CO_2_. The mouse melanoma cell line (B16) was cultured in RPMI 1640 medium (HyClone, USA) supplemented with 10% FBS and 1% penicillin–streptomycin at 37°C and 5% CO_2_. Human OSCC cell lines (UM1, HSC-3, and HN12) and a human renal epithelial cell line (293T) were obtained from the State Key Laboratory of Oral Diseases and maintained in DMEM (Sigma, USA) supplemented with 10% fetal bovine serum (HyClone, USA) at 37°C and 5% CO_2_, and the medium was changed every two days. The cells were passaged at 90% confluence and seeded at 30-40% confluence for maintenance of optimal proliferation conditions.

***Cell transfection and infection***

Six-well plates were used to culture UM1, HSC-3, HN12 and 293T cells, and transfection was carried out when the cells were 70-80% confluent. Serum-free medium was used before transfection reagents were added to the corresponding plates. siRNAs were transiently transfected with Lipofectamine RNAiMAX Transfection Reagent (Thermo Fisher Scientific, USA), and the plasmids were transiently transfected with Lipofectamine 2000 (Invitrogen, USA). After being cultured for 48-72 hours, the cells were seeded. siRNAs for C1QBP were purchased from RiboBio (Guangzhou, China).

***Plasmids***

Plasmids (pcDNA3.1-Myc, pcDNA3.1-Flag, pcDNA3.1-Flag-PA28γ, pcDNA3.1-Myc-PA28γ, pcDNA3.1-Flag-C1QBP, pcDNA3.1-Flag-C1QBP-T76A and pcDNA3.1-Flag-C1QBP-G78N) and C1QBP-related plasmids were synthesized by Genewiz (Su Zhou, China). The full-length C1QBP plasmid was named C1QBP-WT or C1QBP-Full. The C1QBP truncation plasmids pcDNA3.1-Flag-C1QBP 1-167, pcDNA3.1-Flag-C1QBP 94-282 and pcDNA3.1-Flag-C1QBP 168-282 had the amino acid 76-93 functional domain or amino acid 168-213 functional domain removed, respectively.

***Stable cell line generation***

The Flag-PA28γ recombinant lentivirus PA28γ-OE and OE-vector were purchased from NeuronBiotech (Shanghai Genechem Co., Ltd.). UM1, HSC-3 and HN12 cells were infected with the PA28γ-OE or OE-vector. Selective culture medium containing puromycin was used to select cells stably expressing PA28γ-OE or vector controls. The expression of PA28γ was detected by Q-PCR and western blot analysis.

***Cell Invasion Assay***

The in vitro invasion capability of the cells was measured using Transwell chambers coated with Matrigel (Corning, USA; 1:3 dilution with serum-free medium) in 24-well cell culture dishes. Matrigel (35 μL) was added to the upper chamber and incubated for 1 hour at 37°C to ensure that the Matrigel was solidified. Cells were seeded in 200 μl of FBS-free cell culture medium at a density of 5×10^4^ cells/ml in the upper chamber, and the lower chamber was filled with 500 μL of culture medium supplemented with 10% FBS, which acted as a chemoattractant. Cell culture dishes with Transwell chambers were then incubated at 37°C and 5% CO_2_ for 24 hours to allow the cells to invade. At the end of the incubation, the cells on the upper side of the Matrigel-coated filter were removed by wiping with a cotton swab. Crystal violet solution was used to stain invaded cells. The cells were counted under an inverted light microscope (Olympus, Japan). Cells in five randomized fields of view at 200× magnification were counted and are presented as the average number of cells per field of view. The experiment was replicated three times.

***Cell Migration Assay***

The assay was similar to the in vitro invasion assay except that no Matrigel was added to the chamber. The cell concentration in the upper chambers was 5×10^5^/mL, and the cells were added to a 200 μL volume. Cells in five randomized fields of view at 200× magnification were counted and are presented as the average number of cells per field of view. The experiment was replicated three times.

***Cell Proliferation Assay***

After 24 hours of cell transfection, the cells were seeded into a 96-well plate at a density of 5000 cells per well. The culture medium was removed after 24 h and 48 h, respectively. The cell proliferation assay reagent CCK-8 (Beyotime, China) was added under dark conditions, and the ratio of CCK-8 reagent to complete culture medium was 1:9. After incubation in the dark for 1.5 hours, the optical density (OD) of each well was measured under 450 nm light.

***Proximity Ligation Assay***

It was detected using Duolink® PLA Kit (sigma-aldrich, USA) according to the manufacturer’s instructions. The antibodies were PA28γ (1:1000, Cell Signaling Technology, USA), C1QBP (1:400, proteintech, China), IgG_M_ (1:200, Beyotime, China) and IgG_R_(1:200, Beyotime, China).

***Cell*** ***Immunofluorescence Assays***

The cells were added to a 24-well plate (1 × 10^5^ cells per well) with a cover glass at the bottom overnight at 37°C. The cells were fixed with 4% paraformaldehyde for 30 min and then treated with 0.5% Triton X-100 solution for 15 min. Then, the samples were incubated with Tomm20 (1:500, BD Biosciences, USA) and C1QBP (1:400, Abcam, USA) antibodies for 16 hours at 4°C. The next day, TRITC (1:500, Invitrogen, USA) and Alexa Fluor 594 (1:500, Invitrogen, USA) conjugated secondary antibodies were added, and the cells were incubated for 45 min. The cell nuclei were stained with DAPI (Beyotime, China) and observed under a fluorescence microscope. For labeling of mitochondria, the mitochondrial fluorescent dye MitoTracker™ Deep Red FM (Invitrogen, USA) was diluted to 25 nM in culture medium. ImageJ was used to measure the area of mitochondria in the fluorescence micrographs. Mitochondrial Network Analysis (MiNA, available at <https://github.com/ScienceToolkit/MiNA>), an ImageJ-based tool, was used to calculate the mean branch length of mitochondria in fluorescence micrographs.

***Transmission electron microscopy***

The cells were fixed in 4% glutaraldehyde (pH=7.4) and stored in a refrigerator at 4°C. The samples were subsequently sent to Chengdu Lilai Biotechnology Co., Ltd., for osmic acid fixation, neutral resin embedding, and preparation of ultrathin sections. The mitochondrial morphology of the cells was observed using transmission electron microscopy.

***Allogeneic Orthotopic Transplantation Tumor Mouse Model***

All the experiments were conducted in accordance with the guidelines outlined in the Principles of Laboratory Animal Care (NIH) and were approved by the Ethics Committee of West China Hospital of Stomatology, Sichuan University (WCHSIRB-D-2022-023).

Six 8-week-old C57BL/6 mice were used as experimental animals and were purchased from Chengdu Dossy Experimental Animals Co., Ltd. Submucous injections into the tongue of each mouse were performed. A total of 1×10^6^ control or PA28γ-overexpressing 4MOSC2 cells were injected into the mice subcutaneously, and the mice were divided into a control group (PA28γ NC, n=3) and an experimental group (PA28γ OE, n=3). Ten days later, carbon dioxide euthanasia was carried out. The collected tongue was cut along the midline of the tumor, and half of the tissue was fixed in 4% paraformaldehyde solution for 48 hours. After dehydration and embedding, subsequent hematoxylin and eosin (HE) staining was carried out, and the other half of the isolated tumor tissue was frozen at -80 ℃ for ATP and ROS analysis.

***Subcutaneous Transplantation Tumor Mouse Model***

A total of 24 C57BL/6 mice aged 6-8 weeks were used as experimental animals, of which 12 were Pa28γ whole-body knockout mice, with the other 12 being the same litter, and same-sex wild-type control mice were obtained from our research group. Submucous injections into the back of each mouse were performed. A total of 1×10^6^ cells (100μL) were injected subcutaneously, and mice were divided into the following 4 groups based on different genotypes and cell lines: Pa28γ wild-type mice injected with control B16 cells (Pa28γ^wt/wt^ + Vector, n=6), Pa28γ wild-type mice injected with PA28γ knockdown B16 cells (Pa28γ^wt/wt^ + shPa28γ, n=6), Pa28γ whole-body knockout mice injected with control B16 cells (Pa28γ^ko/ko^ + Vector, n=6) and Pa28γ whole-body knockout mice injected with PA28γ knockdown B16 cells (Pa28γ^ko/ko^ + shPa28γ, n=6). On the 7th day after injection of tumor cells, the formation of subcutaneous tumors in the back was visible. On the 15th day, the mice were euthanized with carbon dioxide, and the back tumor tissue was collected. The tumor volume was calculated as (length × width^2^)/2 after excision. Each tumor was half placed in 4% paraformaldehyde to prepare paraffin sections and half stored in the -80 °C for other the ATP and ROS analysis.

***Mitochondrial function detection***

Mitochondrial OXPHOS function was detected using a Seahorse XFe24 Analyzer (Agilent, USA) and Seahorse XF Cell Mito Stress Test Kit (Agilent, USA) according to the manufacturer’s instructions. Cells were seeded in an XF24-well cell culture plate at a density of 10000 cells/well for 24 h at 37°C and 5% CO2. The medium was changed to 500 µL of Seahorse XF DMEM (pH 7.4), and the cells were incubated at 37°C without 1.5 µM oligomycin, 2 µM carbonyl cyanide-4-(trifluoromethoxy) phenylhydrazone (FCCP) or 0.5 µM rotenone and antimycin A to test ATP-linked respiration, the maximal OCR and nonmitochondrial oxygen consumption, respectively. All OCRs were normalized according to the cell number.

For ROS production, cells were seeded into a 96-well plate (1 × 10^4^ cells per well) overnight in the dark at 37°C. Then, 100 µL of ROS dye-containing culture medium [the ROS fluorescent probe DHE (KeyGEN, China) at a concentration of 100 µM] was added to each well, and the mixture was incubated for 45 minutes. After the cells were washed twice with PBS, the fluorescence intensity was measured at an excitation wavelength of 518 nm and an emission wavelength of 605 nm. ROS levels in tumors were calculated using a ROS detection kit (Beyotime, China) following the manufacturer’s protocol. Ten milligrams of tissue from each group were cut, and the tumors in each group were placed in the same tube. Then, the tumors were homogenized into tissue homogenates using a homogenizer at 4 ℃ in 1 mL of lysis solution, after which the protein concentration was determined via the BCA method after ultrasonic lysis. Solutions for blank, standard, control and test tubes were prepared according to the protocol. The tubes were kept at 37°C for 2 min, after which 2 mL of developer was added. The solutions were mixed well and left at room temperature for 20 minutes before being measured at 550 nm in a microplate reader.

The ATP production of tumors was calculated using an ATP detection kit (KeyGEN, China) according to the manufacturer’s protocol. Ten milligrams of tissue homogenate from each tumor were prepared using a homogenizer at 4°C in 100 µL of lysis solution, after which the protein concentration was determined via the BCA method after ultrasonic lysis. An ATP standard solution was prepared with lysis solution to construct standard curves at concentrations of 0.01, 0.03, 0.1, 0.3, 1, 3, and 10 µM. Then, 100 µL of detection solution was added to 96-well plates and kept at room temperature for 5 minutes. Then, 20 µL of standard sample or tumor sample was added to the wells. The spontaneous fluorescence intensity of each well was tested, and ATP production was calculated according to standard curves.

***RNA Extraction and RT‒qPCR Analysis***

Total RNA was extracted using TRIzol Reagent (Invitrogen, USA) according to the manufacturer’s instructions. The primers for PSME3, C1QBP and β-actin were purchased from RiboBio (Guangzhou, China). A FastStart Universal SYBR-Green Master Mix kit was purchased from Roche (Mannheim, Germany). qPCR for the detection of PSME3 and C1QBP mRNA expression levels was performed in an ABI 7500 Real-Time PCR system (ABI, Foster City, USA). Relative mRNA expression was normalized to that of β-actin. The specific primer sequences for the genes used were as follows:

β-actin forward: 5’- CACCATTGGCAATGAGCGGTTC -3’,

reverse: 5’- AGGTCTTTGCGGATGTCCACGT -3’;

PSME3 forward: 5’- AAGGTTGATTCTTTCAGGGAGC -3’,

reverse: 5’-AGTGGATCTGAGTTAGGTCATGG -3’;

C1QBP forward: 5’- CACACCGACGGAGACAAAG -3’,

reverse: 5’- GGGAGGGTTTTATGCTTCTGAAT -3’;

***Western Blot Assay***

The cells were lysed in lysis buffer (1% NP-40 supplemented with a complete protease inhibitor tablet (self-formulated)) on ice. Protein extracts (30 μg) were separated by a 10% SDS‒PAGE gel and transferred to 0.45 μm polyvinylidene difluoride (PVDF) membranes (Millipore, USA). After blocking with 5% bovine serum albumin (BSA) for 1 hour, the membranes were incubated with the following primary antibodies overnight at 4°C: α-Tubulin (Cell Signaling Technology, USA, 1:5000 dilution), Flag (Cell Signaling Technology, USA, 1:2000 dilution), Myc (Cell Signaling Technology, USA, 1:1000 dilution), C1QBP (Cell Signaling Technology, USA, 1:1000 dilution), PA28γ (Cell Signaling Technology, USA, 1:1000 dilution), OPA1 (Cell Signaling Technology, USA, 1:1000 dilution), Mitofusin-1 (Cell Signaling Technology, USA, 1:1000 dilution), Mitofusin-2 (Cell Signaling Technology, USA, 1:1000 dilution) and Total OXPHOS Rodent WB Antibody Cocktail (Abcam, USA, 1:1000 dilution). Afterwards, the membranes were incubated with either an anti-goat, anti-mouse or anti-rabbit HRP-conjugated secondary antibody (Sigma–Aldrich, Australia, 1:3000 dilution) for 1 h. After incubation with a chemiluminescence substrate, images were taken with an ImageReader LAS-2000 (Fujifilm, Japan), and the proteins were analyzed with ImageJ software.

***Immunoprecipitation assay***

The cells were lysed in lysis buffer (1% NP-40 supplemented with a complete protease inhibitor tablet (self-formulated) on ice, and 20 μL of cell lysate was used as the input. The cell lysates were coincubated with antibodies at 4°C overnight. The protein G agarose was washed and added to a 1 mL sample at a ratio of 20 µl. The sample was shaken on a horizontal shaker for 4 h, after which the deposit was collected for western blotting.

***HE Staining Analysis***

The tissues were immersed in 4% paraformaldehyde for 24 hours and washed overnight with running water. The tissue was then placed in an automatic dehydrator. After dehydration, the tissue was embedded in paraffin and sectioned at a thickness of 5 μm. Dewaxing and gradient alcohol hydration were conducted before the sections were counterstained with hematoxylin and eosin. After dehydration with an alcohol gradient, the sections were sealed with neutral gum.

***Immunohistochemistry*** ***and Analysis***

Immunohistochemistry (IHC) for PA28γ (Cell Signaling Technology, USA, 1:400 dilution) and C1QBP (Abcam, USA, 1:800 dilution) was performed on FFPE specimens after antigen retrieval with EDTA buffer (1 mM, pH 8.0), and the proteins were visualized by diaminobenzidine (DAB, Gene Tech, 1:50 dilution). The MVD was calculated as the average count of microvessels in four hot spots at high magnification (400×). The intensity of staining (0, no staining; 1, weakly stained; 2, moderately stained; 3, strongly stained) was assessed by an experienced pathologist without any knowledge of the clinical or pathological data, and the percentage of positive cells was noted. The staining proportions were scored as 0 (≤5%), 1 (5-33%), 2 (34-66%), or 3 (≥67%), and the total staining was expressed as a product of the two numbers (six levels: 1, 2, 3, 4, 6 and 9). For the statistical analysis of prognostic value in the OSCC cohort, the total staining scale was divided into two categories: 1 (staining scale ≤4) and 2 (staining scale >4).

***Bioinformatics***

To identify the specific genes targeted by PSME3 and C1QBP in OSCC, GeneMANIA was used. To determine the prognostic value of C1QBP, Gene Expression Profiling Interactive Analysis (GEPIA) (http://gepia.cancer-pku.cn) was used to analyze the survival of OSCC patients in the TCGA HNSC and SKCM database.

***Prediction and Analysis of Protein Interactions***

**Protein Sequence Retrieval and Structure Prediction**

The protein sequences of C1QBP and PA28γ were obtained from the AlphaFold Protein Structure Database. Structural predictions of the protein-protein interaction between C1QBP and PA28γ were conducted using AlphaFold 3. The plDDT (predicted local distance difference test) values were utilized to assess the confidence of the predicted models. Models with a plDDT score above 70 were considered confident, while those with a score above 90 were categorized as very high confidence. These values were annotated in the figures to indicate the reliability of the structural predictions.

**Protein Preparation and Structure Optimization**

The best-scored model for the C1QBP-PA28γ interaction predicted by AlphaFold 3 was selected for further analysis. The model was imported into MOE 2022 (Molecular Operating Environment) software for protein preparation. This process included the removal of water molecules and other heteroatoms, followed by the addition of hydrogen atoms to the structure. This step was essential for optimizing the protein’s 3D conformation and ensuring the correctness of the protonation states at physiological pH.

**Energy Minimization and Hydrogen Bond Prediction**

The protein structure was subjected to energy minimization using the Amber10: EHT (Effective Hamiltonian Theory) force field, with R-field 1: 80 settings to refine the model’s geometry. The minimization process was performed to optimize the protein’s internal energy and ensure stable conformation, followed by calculation of hydrogen bond interactions. The interaction energies and hydrogen bonds were analyzed to identify potential binding sites and stabilize the predicted protein-protein complex.

**Appendix figures and figure legends**

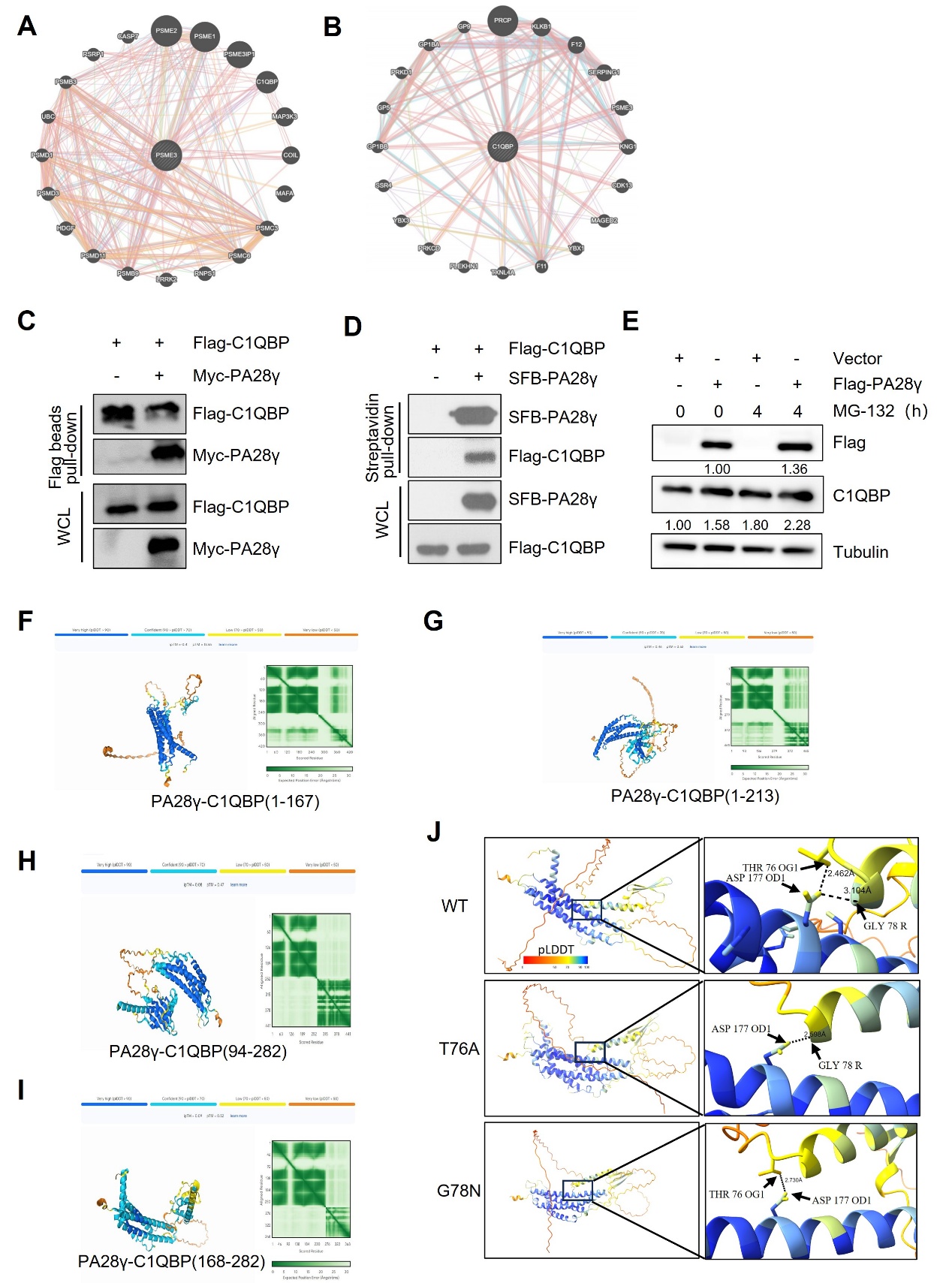

**Appendix Fig. 1** **The interaction between PA28γ and C1QBP. (A, B)** The gene network of PA28γ and C1QBP in the GeneMANIA database. **(C, D)** The interaction between exogenous PA28γ and C1QBP in 293T cells was verified via pull-down. **(E)** 293T cells transfected with Vector or Flag-PA28γ were treated with MG-132 (10μM) for the indicated periods of time. **(F-I)** The prediction of interactions between PA28γ and truncated C1QBP with AlphaFold 3. **(J)** The prediction of interactions between PA28γ and C1QBP wild type or mutations with AlphaFold 3.

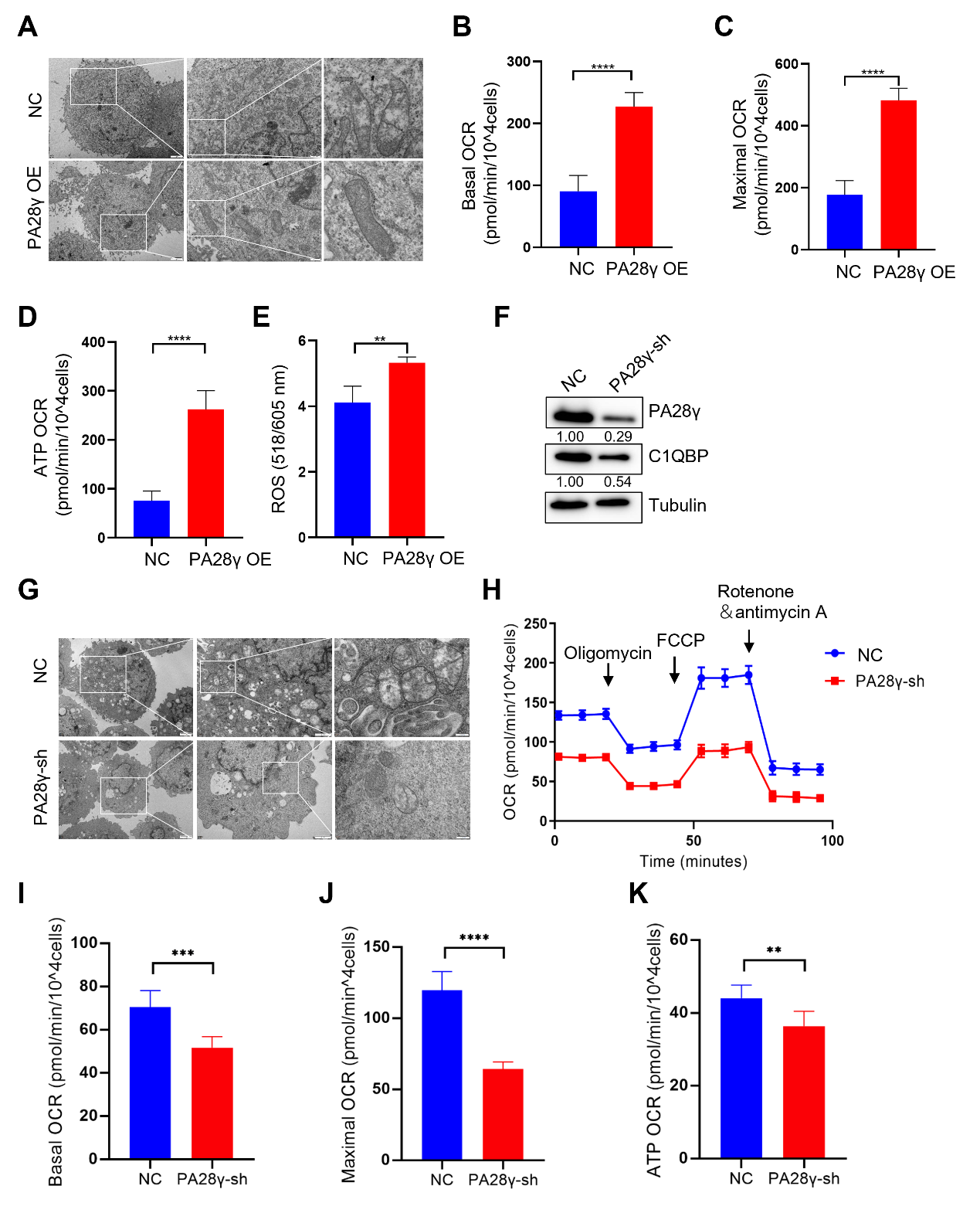

**Appendix Fig. 2** **PA28γ and C1QBP colocalize in mitochondria and regulate OXPHOS in OSCC cells**. **(A)** TEM images of PA28γ-overexpressing and control HN12 cells. **(B-D)** Basal OCRs, maximal OCRs and ATP production of PA28γ-overexpressing and control HN12 cells measured by the cell Mito Stress Test (the data are presented as the means ± SDs of 3 independent experiments; *****P*<0.0001). **(E)** ROS generation in PA28γ-overexpressing and control HN12 cells (the data are presented as the means ± SDs of 3 independent experiments; ***P*<0.01). **(F)** Western blot analysis of PA28γ-sh and control UM1 cells. **(G)** TEM images of PA28γ-sh and control UM1 cells. **(H)** OCRs of PA28γ-sh and control UM1 cells were plotted using a Cell Mito Stress Test Kit (the data are presented as the means ± SDs of 3 independent experiments). **(I-K)** Basal OCRs, maximal OCRs and ATP production of PA28γ-sh and control UM1 cells measured by the Cell Mito Stress Test (the data are presented as the means ± SDs of 3 independent experiments; ***P*<0.01, ****P*<0.001, and *****P*<0.0001).

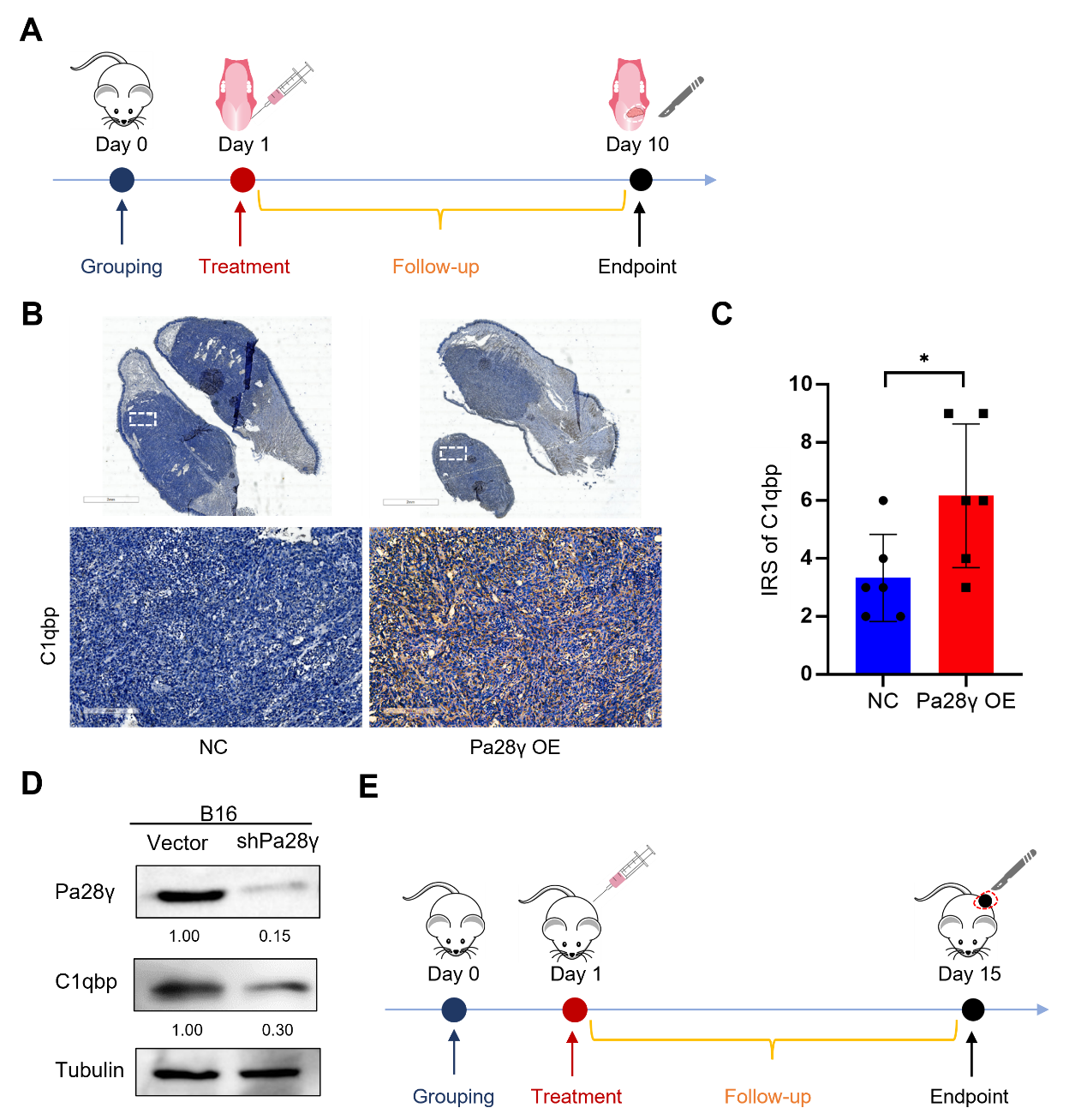

**Appendix Fig. 3 PA28γ overexpression can upregulate C1QBP in vivo. (A)** Diagram of allogeneic orthotopic transplantation tumor mouse model. **(B)** Representative IHC images of C1QBP antibody staining in Pa28γ-overexpressing and control nude mice. **(C)** IRSs of C1QBP antibody staining in the Pa28γ-overexpressing and control nude mice (n=6, the data are presented as the means ± SDs; **P*<0.05). **(D)** Western blot analysis of PA28γ-silenced and control B16 cells. **(E)** Diagram of subcutaneous transplantation tumor mouse model.

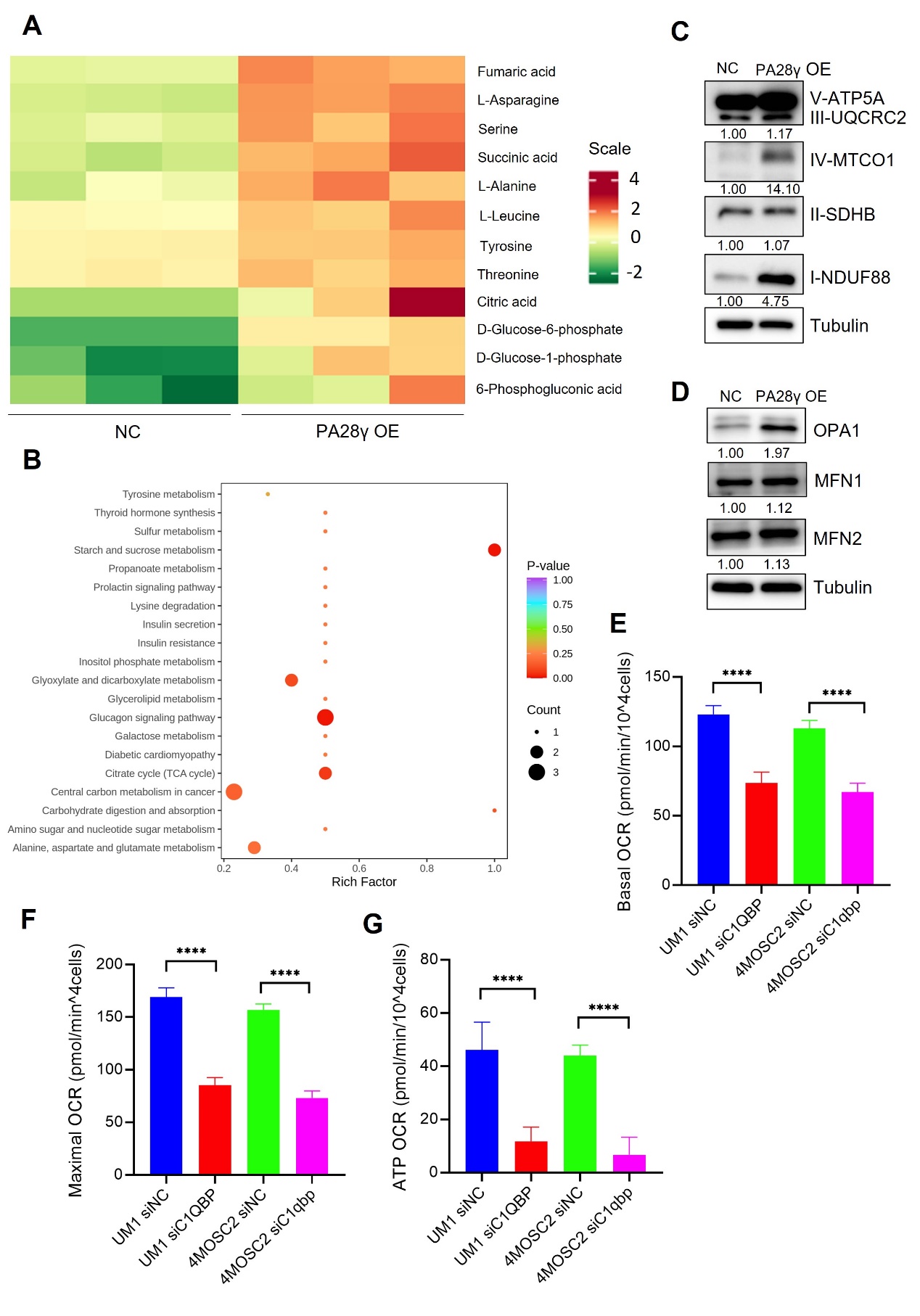

**Appendix Fig. 4 PA28γ and C1QBP are involved in OXPHOS and cellular biological behavior. (A, B)** Heatmap differential metabolites and KEGG enrichment analysis in the PA28γ-overexpressing and control HN12 cells. **(C, D)** Western blot analysis of PA28γ-overexpressing and control HN12 cells. **(E-G)** Basal OCRs, maximal OCRs and ATP production of C1QBP-silenced and control PA28γ-overexpressing UM1 and 4MOSC2 cells measured by the Cell Mito Stress Test (data are presented as the mean ± SD of 3 independent experiments, *****P*<0.0001).

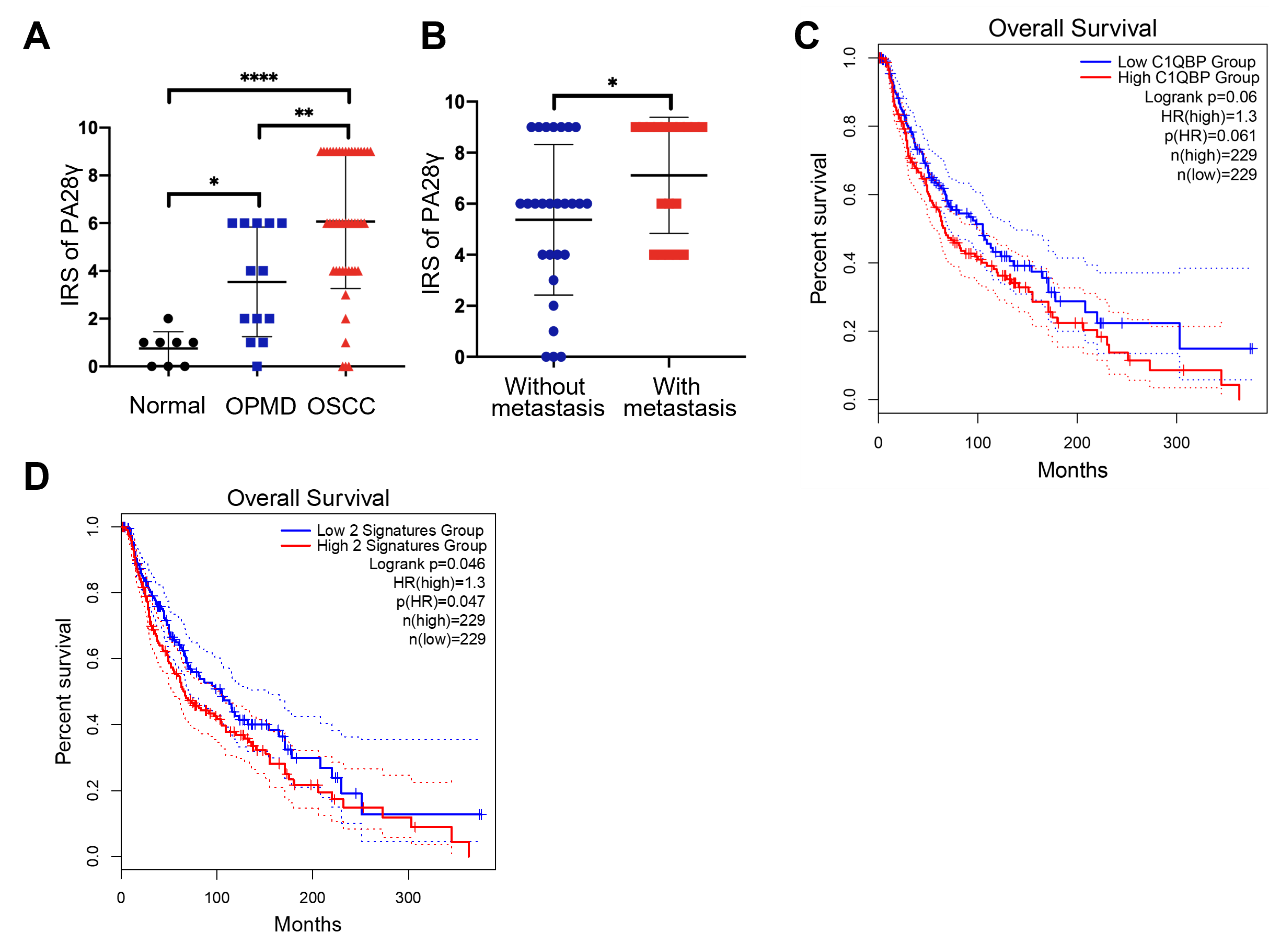

**Appendix Fig. 5 PA28γ and C1QBP are involved in the development of tumor.** **(A)** Comparison of the IRSs of PA28γ between the normal (n=8), OPMD (n=13) and OSCC (n=45) groups (the data are presented as the means ± SDs; **P*<0.05, ***P*<0.01, *****P*<0.0001). **(B)** Comparison of the IRSs of PA28γ in the nonmetastatic (n=27) and metastatic (n=18) OSCC groups (the data are presented as the means ± SDs; **P*<0.05). **(C)** Kaplan–Meier analysis of the protein expression of C1QBP in TCGA SKCM database (n=229, *P*=0.061). **(D)** Kaplan–Meier analysis of both low or high protein expression of C1QBP and PA28γ in TCGA SKCM database (n=229, *P*=0.047).

**Appendix Table 1. Baseline Characteristics of the Patients with Oral Squamous Cell Carcinoma in Cohort I**

| **characteristic** |  | N=45  *Number (%)* | *P* value |
| --- | --- | --- | --- |
| Gender |  |  |  |
| male |  | 26（57.8） | 0.7553 |
| female |  | 19（42.2） |  |
| Age（year） |  |  |  |
| ﹤60 |  | 11（24.4） | 0.3428 |
| ≥60 |  | 34（75.6） |  |
| Tumor stage |  |  |  |
| T1-2 |  | 30（66.7） | 0.0030 |
| T3-4 |  | 15（33.3） |  |
| Node stage |  |  | 0.0217 |
| N0 |  | 27（60.0） |  |
| N1-3 |  | 18（40.0） |  |
| Clinical stage |  |  |  |
| I-II |  | 23（51.1） | 0.0045 |
| III-IV |  | 22（48.9） |  |
| Differentiation degree |  |  | 0.2651 |
| Low/Moderate |  | 11（24.4） |  |
| High |  | 34（75.6） |  |
| Recurrence |  |  |  |
| Yes |  | 26（57.8） | 0.5900 |
| No |  | 19（42.2） |  |
| C1QBP expression |  |  |  |
| low |  | 13（28.9） | ＜0.05 |
| high |  | 32（71.1） |  |

**Appendix Table 2. Baseline Characteristics of the Patients with Oral Squamous Cell Carcinoma in Cohort II**

| **characteristic** |  | N=295  *Number (%)* | *P* value |
| --- | --- | --- | --- |
| Gender |  |  |  |
| male |  | 216（73.2） | 0.0798 |
| female |  | 79（26.8） |  |
| Age（year） |  |  |  |
| ﹤60 |  | 126（42.7） | 0.6825 |
| ≥60 |  | 169（57.3） |  |
| Smoking |  |  |  |
| Yes |  | 145（49.2） | 0.9483 |
| No |  | 150（50.8） |  |
| Tumor stage |  |  |  |
| T1-2 |  | 219（74.2） | 0.7181 |
| T3-4 |  | 76（25.8） |  |
| Node stage |  |  | 0.5265 |
| N0 |  | 194（65.8） |  |
| N1-3 |  | 101（34.2） |  |
| Clinical stage |  |  |  |
| I-II |  | 155（52.5） | 0.9280 |
| III-IV |  | 140（47.5） |  |
| Recurrence |  |  |  |
| Yes |  | 149（50.5） | 0.1889 |
| No |  | 146（49.5） |  |
| C1QBP expression |  |  |  |
| low |  | 132（44.7） | ＜0.05 |
| high |  | 163（55.3） |  |

**Appendix Table 3. Baseline Characteristics of the Patients with Head and Neck Squamous Cell Carcinoma in TCGA cohort**

| **Characteristic** | **C1QBP expression** | | *P* value |
| --- | --- | --- | --- |
|  | Low (N=259, *Number* %) | High (N=259, *Number* %) |  |
| Gender |  |  |  |
| male | 190（73.3） | 193（74.5） | 0.8414 |
| female | 69（26.7） | 66（25.5） |  |
| Age（year） |  |  |  |
| ﹤60 | 133（51.4） | 100（38.6） | 0.0047 |
| ≥60 | 126（48.6） | 159（61.4） |  |
| Tumor stage* |  |  |  |
| I-II | 57（22.0） | 40（15.4） | 0.0384 |
| III-IV | 161（62.2） | 186（71.8） |  |
| Clinical stage* |  |  |  |
| I-II | 66（25.5） | 51（19.7） | 0.1702 |
| III-IV | 189（73.0） | 199（76.8） |  |
| Histological grade* |  |  |  |
| G1-2 | 174（67.2） | 191（73.7） | 0.1876 |
| G3-4 | 72（27.8） | 60（23.2） |  |

* The information of some patients is missing.

**Appendix Table 4. Baseline Characteristics of the Patients with Skin Cutaneous Melanoma in TCGA cohort**

| **Characteristic** | **C1QBP expression** | | *P* value |
| --- | --- | --- | --- |
|  | Low (N=229, *Number* %) | High (N=229, *Number* %) |  |
| Gender |  |  |  |
| male | 143（62.4） | 145（63.3） | 0.9230 |
| female | 86（37.6） | 84（36.7） |  |
| Age（year） |  |  |  |
| ﹤60 | 107（46.7） | 102（44.5） | 0.7075 |
| ≥60 | 122（53.3） | 127（55.5） |  |
| Tumor stage* |  |  |  |
| I-II | 77（33.6） | 71（31.0） | 0.6013 |
| III-IV | 117（51.1） | 122（53.3） |  |
| Node stage* |  |  |  |
| 0 | 106（46.3） | 123（53.7） | 0.1599 |
| 1-3 | 94（41.0） | 81（35.4） |  |
| Metastasis stage* |  |  |  |
| 0 | 205（89.5） | 204（89.1） | 0.9999 |
| 1 | 11（0.05） | 11（0.05） |  |
| Clinical stage* |  |  |  |
| I-II | 102（44.5） | 118（51.5） | 0.1373 |
| III-IV | 103（45.0） | 87（38.0） |  |

* The information of some patients is missing.
